# Reagent- and actuator-free analysis of individual erythrocytes using three-dimensional quantitative phase imaging and capillary microfluidics

**DOI:** 10.1101/2021.05.15.442583

**Authors:** DongHun Ryu, Hyeono Nam, Jessie S. Jeon, YongKeun Park

## Abstract

Histopathological examination of blood cells plays a crucial role in the diagnosis of various diseases. However, it involves time-consuming and laborious staining procedures required for microscopic review by medical experts and is not directly applicable for point-of-care diagnosis in resource-limited locations. This study reports a dilution-, actuation- and label-free method for the analysis of individual red blood cells (RBCs) using a capillary microfluidic device and quantitative phase imaging. Blood, without any sample treatment, is directly loaded into a micrometer-thick channel such that it forms a quasi-monolayer inside the channel. The morphological and biochemical properties of RBCs, including hemoglobin concentration, hemoglobin content, and corpuscular volume, were retrieved using the refractive index tomograms of individual RBCs measured using 3D quantitative phase imaging. The deformability of individual RBCs was also obtained by measuring the dynamic membrane fluctuations. The proposed framework applies to other imaging modalities and biomedical applications, facilitating rapid and cost-effective diagnosis and prognosis of diseases.

## Introduction

The monitoring and assessment of the morphology and biochemistry of blood cells are the most fundamental laboratory analyses for screening and diagnosing various diseases, such as sepsis^1, 2^ and leukemia^2^. Complete blood count (CBC) tests based on absorption spectroscopy, impedance measurement, and fluorescence flow cytometry are used routinely at wet labs and clinical sites^3,4^. While these standard techniques assess important hematological parameters, such as hemoglobin (Hb) concentration, mean cellular volume, and differential counts of several white blood cells, they are complex and expensive^5^. Furthermore, additional microscopic review of a stained blood smear must often be performed manually by medical professionals to handle any flagged specimens, necessitating a significant amount of sample treatment that requires time and labor^6^.

Label-free methods for imaging and characterization of blood cells, such as Raman microscopy^7^ and quantitative phase imaging (QPI)^8–13^, have been extensively investigated to circumvent the abovementioned issues. These methods exploit the intrinsic contrast of a blood cell to probe its morphological and biochemical properties without using any labeling agents, simplifying the sample preparation steps significantly. Nonetheless, a simple yet critical impediment to a rapid point-of-care (POC) hematological test has remained unaddressed in previous approaches wherein the sample concentration must be controlled manually for optimal imaging. To fully analyze individual blood cells, the cells need to be prepared with optimal concentrations, both laterally and axially, for 3D imaging. Additionally, blood cells in a suspension must be sufficiently static while being captured. The steps associated with practical constraint in blood cell imaging require additional time-consuming and laborious troubleshooting processes.

In recent years, microfluidic technologies have been exploited to simplify such sample preparation steps, enabling precise cell control and separation^14–16^. While conventional microfluidic controllers and separators with external pumps, exploiting electric ^17, 18^, magnetic^19, 20^, or acoustic forces^21, 22^, have played a major role in diverse POC tests, recent microfluidic biosensors that do not require any external actuators have also been explored using finger actuation^23, 24^, capillary force^25, 26^, paper substrates^24, 27^, degassed systems^28, 29^, and gas-generating systems^30, 31^. Among these, capillary-force-based microfluidic devices have been considered user-friendly and straightforward. Additionally, the device does not require any external actuators. Its fluid properties, such as the sample filling rate, can be directly determined using the device’s dimensions and material properties^32, 33^, enabling consistent and straightforward POC tests.

This study presents a dilution-, actuation- and label-free framework for the characterizing of morphological, biochemical, and mechanical properties of individual red blood cells (RBCs) using refractive index (RI) tomography and capillary microfluidic devices to enable rapid and cost-effective blood tests (Fig. 1). Using a SiO2 microfluidic device, which is capable of generating a quasi-monolayer of blood cells through capillary action from the whole blood, we performed 3D QPI imaging of RBCs and subsequently obtained important parameters of RBCs, such as Hb concentration, content, and cellular volume analyzing the reconstructed 3D RI tomogram of individual cells^34, 35^. Furthermore, the approach allowed us to study the mechanical deformability of RBCs, which has been used to reveal the etiology of human diseases^36–39^. We acquired dynamic topographic maps of RBCs, stably captured by our device, and quantified their membrane fluctuation values using the maps. We also explored different device dimensions and quantified the extent to which the blood cells filled and overlapped in the microchannel without any actuators to provide a reference point for optimal device specifications. We believe that the proposed blood-testing platform, which does not require expensive and destructive labeling reagents and instruments, will enable various POC applications beyond those presented in this work.

**Figure 1.**
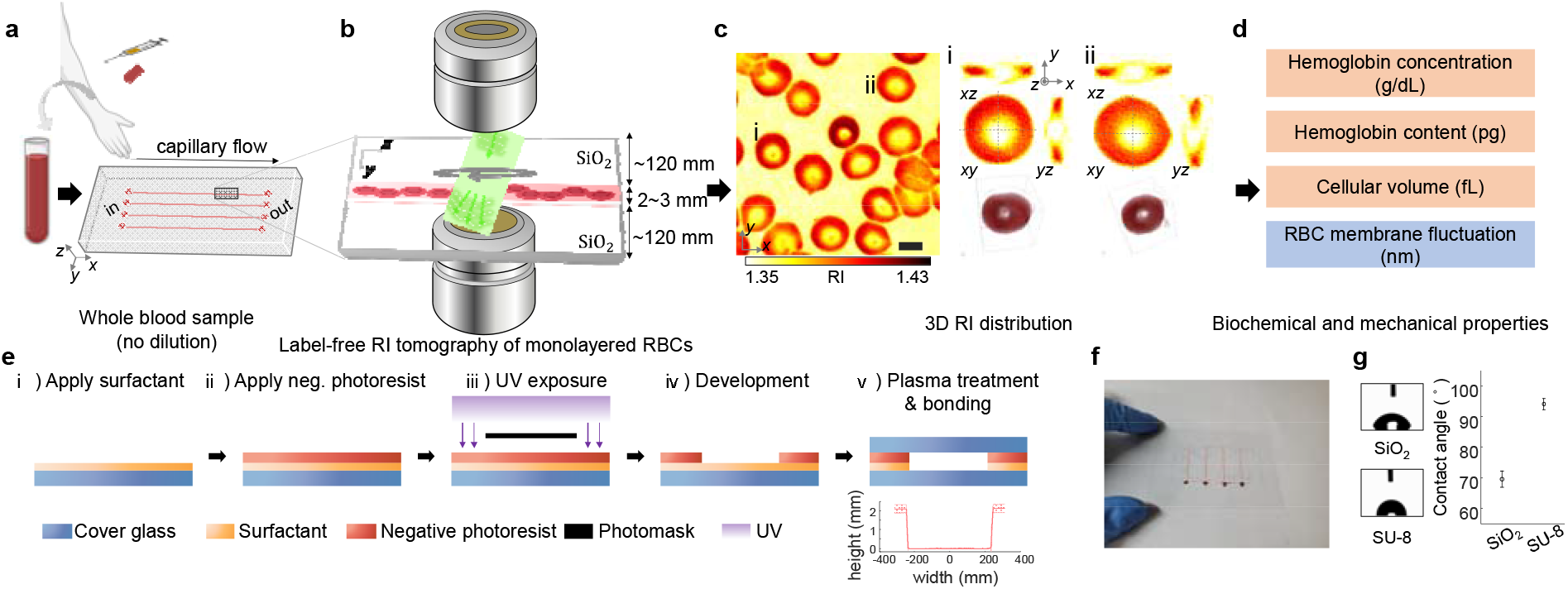
Overview of our dilution-free, label-free RBC test framework. (a) Whole blood cells loaded into the capillary microfluidic device without dilution or treatment. (b) Quasi-monolayer RBCs inside the microfluidic channel for label-free refractive index (RI) tomographic imaging. (c) 3D RI tomograms of RBCs. Two representative RBCs visualized from three different perspectives and their iso-surface renderings. (d) Biochemical and mechanical properties, including Hb concentration, Hb content, cellular volume, and membrane fluctuation, obtained directly from the tomographic imaging (See Methods). (e) Fabrication of the microfluidic device based on negative photolithography. (f) Device photograph. (g) Channel height measured via a surface profiler (mean: 2.12 μm, std: 0.23 μm). (h) Contact angle of the materials constituting the cross-section of the channel. Negative capillary pressure due to low contact angle (<90°) allows the sample to be loaded into the channel withoutexternal forces. Scale bar = 5 μm.

## Methods

### Microfluidic device fabrication

The capillary flow-driven microfluidic device developed in this study, which enables dilution-free RBC imaging, was fabricated based on photolithography (Fig. 1e). A 50 × 70 mm^2^ SiO_2_ microscope cover glass (Matsunami, Japan) was used for the bottom of the device. A surfactant (OmniCoat, Kayaku Advanced Materials, USA) was applied following the manufacturer’s protocol to increase the adhesion between the glass and the negative photoresist. The SU-8 2005 or 2007 negative photoresist (SU-8 2000, Kayaku Advanced Materials, USA) was then spin-coated to match the target height of the device. After soft-baking it on a hot plate, the cover glass coated with SU-8 was subjected to UV exposure to create the desired height microchannel. The device substrate was then bonded with a 24×50 mm^2^ SiO_2_ microscope cover glass (Thickness No. 1, Matsunami, Japan) via plasma treatment. In this study, 2, 5, and 7 μm devices were fabricated. The 2 μm device was used to obtain the main results from the characterization of various RBC properties.

### Channel height measurement

The device channel height was measured using a surface profiler (Dektak-8, VEECO, USA), as shown in Fig. 1g. As the surface profiler is in physical contact with the device surface to profile its height, no cover glass was bonded over the negative photoresist for the devices used to measure the channel height.

### Contact angle measurement

The contact angles of the materials constituting the device were measured using a contact angle analyzer (Phoenix 300 Plus, SEO Co., Korea). A 3 μL water droplet was placed on the surface of each material, and the built-in software was used to calculate the contact angle. The key difference between the conventional devices and the one developed in this study is the use of SiO_2_ to achieve the capillary flow of blood cells inside the channel. The low contact angle achieved using our device enables the fluid to flow autonomously without any external forces.

### Channel flow measurement

The time required for the liquid to fill the device and its flow front position were measured to test the fluid dynamics inside our device. Instead of the whole blood, the viscosity of which changes over time, red-dyed distilled water was dropped through the inlet. The time required to reach the outlet (24 mm) was measured and used to calculate the filling rate. The Fiji software was used to measure the flow front position (https://imagej.net/Fiji). Finally, the experimental flow front over time was compared to a theoretical model (for more details, see also Supplementary Information) that considers liquid properties such as density, dynamic viscosity, surface tension coefficient, and channel specifications.

### Imaging system and reconstruction procedure

For 3D QPI imaging, we used a commercialized optical diffraction tomography (ODT) microscope (also known as holotomography), based on Mach–Zehnder interferometry, equipped with a digital micromirror device (DMD) for angle-scanning illumination (HT-2H, Tomocube, Inc., South Korea) to image the blood cells inside our device^40^. A schematic of the optical setup is shown in Fig. 2a. The interferometric setup utilizes a diode-pumped solid-state laser beam with the wavelength λ = 532 nm and splits the input beam into a sample beam and reference beam via a fiber coupler. The sample beam was angle-modulated by a DMD (DLP65300FYE, Texas Instruments, USA) and impinged on the blood cells inside the microfluidic device through an objective lens (UPLASAPO 60XW, Olympus Inc., Japan). The scattered light, 4-*f*-relayed by an objective lens (UPLASAPO 60XW, Olympus Inc., Japan) and a tube lens, interfered with the reference beam transmitted by a beam splitter to form an interferogram at the camera plane after being filtered by a linear polarizer.

**Figure 2.**
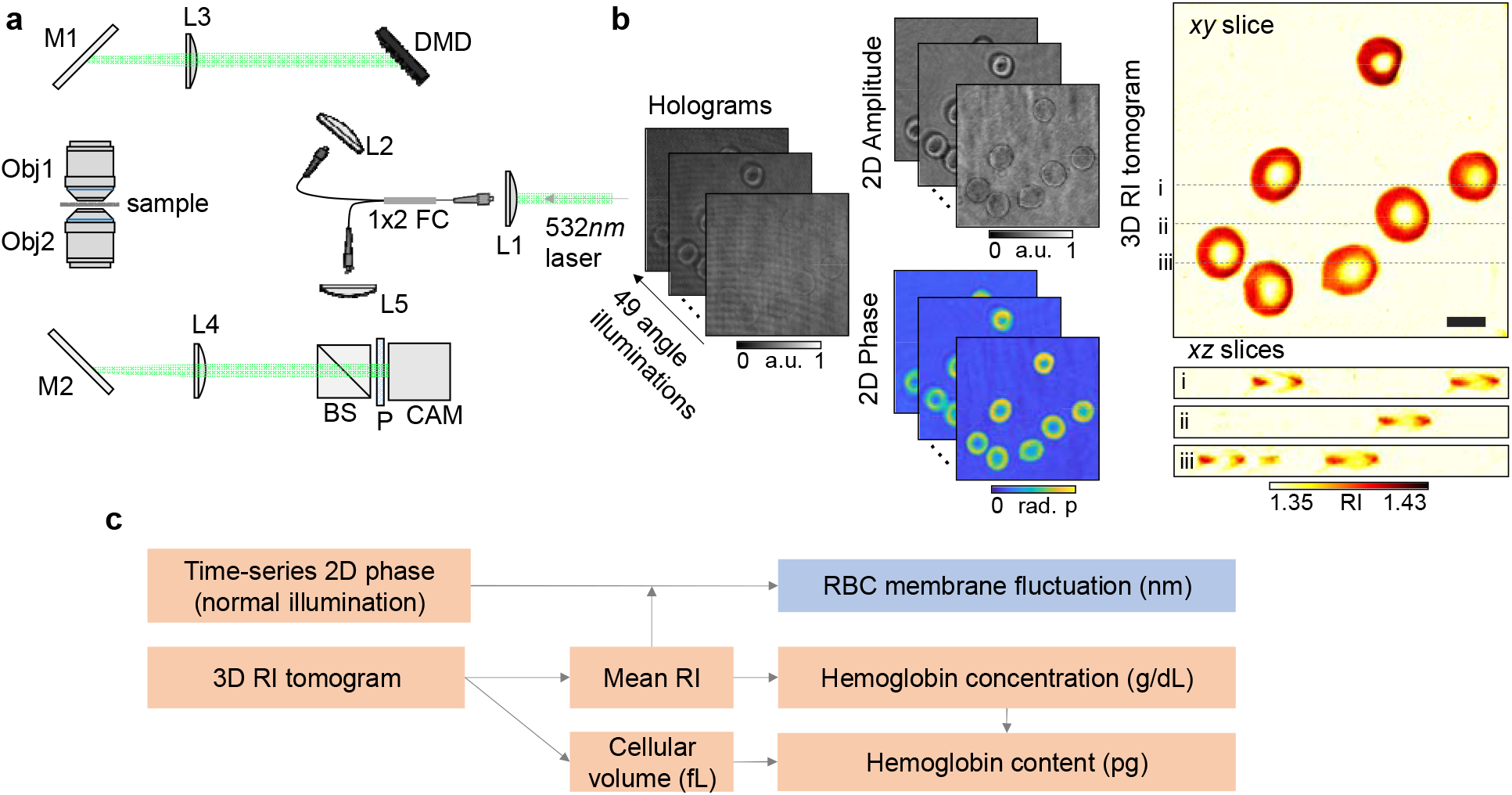
Tomographic reconstruction and RBC parameter quantification. (a) Schematics of the optical setup. (b) Tomographic reconstruction process. Complex field information retrieved from the angle-scanned holograms is processed to reconstruct a 3D refractive index tomogram. (c) Four quantitative parameters obtained from the RI tomogram and time-series phase delay map. L: Lens. FC: Fiber coupler. DMD: Digital micromirror device. M: Mirror. Obj: Objective lens. BS: Beam splitter. P: Polarizer. CAM: Camera. Scale bar = 5 μm.

Forty-nine interferograms were obtained and processed to retrieve each complex field image using a spatial-filtering-based phase-retrieval algorithm z and the Goldstein phase unwrapping method to reconstruct the RI tomograms of RBCs, as depicted in Fig. 2b^41^. A 3D RI map was then reconstructed from the complex field images based on the Fourier diffraction theorem with the Rytov approximation^42–44^. The 3D RI distribution of a sample was reconstructed from the retrieved amplitude and phase images. A non-negativity regularization algorithm was applied to the reconstructed 3D tomogram to overcome the missing cone problem due to the limited numerical apertures of the objective lenses ^45^.The theoretical resolution of our imaging system was 110◻nm (lateral) and 330 nm (axial) according to the Lauer criterion ^46, 47^. The detailed principles and the reconstruction algorithms of the tomographic reconstruction can be found elsewhere^48, 49^.

### Quantification of RBC parameters

Figure 2c shows the process of obtaining the quantitative parameters of RBCs. Hemoglobin (Hb) concentration, content, and cellular volume were obtained from the reconstructed 3D RI tomogram. Cytoplasmic Hb concentration was computed by exploiting the linear relationship between the RI of the Hb solution and its Hb concentration, assuming that RBC cytoplasm is mainly consisted with HB proteins, as follows: <n – n_0_> = α [Hb], where n and n_0_ are the RI values of the voxel and medium, respectively, and α is the refractive index increment (RII)^50, 51^. Setting α = 0.18 mL/g and n_0_ = 1.35, measured using a refractometer (Atago™ R-5000, Japan), the Hb content was directly obtained from the Hb concentration using the cellular volume computed by thresholding the tomogram.

The membrane fluctuation of the RBC was obtained from time-series phase measurements^9, 52, 53^. As the membrane height of an RBC is directly related to its optical phase delay under normal illumination, we quantified the membrane fluctuation by computing the temporal change in the RBC height profiles h(x,y,t) converted from the optical phase delay Δø, as Δø = (2π/λ)Δn· h, where Δn is an RI difference between a RBC and a surrounding medium. The 2D membrane fluctuation map, σ_h(x,y)_, is defined as the temporal standard deviation of the RBC height profiles, as follows: σ_h(x,y)_ = [<(h(x,y,t) - <h(x,y,t)>_time_)^2^>_time_]1/2, and the membrane fluctuation quantity σ_h_ is a spatially averaged value of σ_h(x,y)_ over the sample area: σ_h_ = < σ_h(x,y)_>_space_.

### Sample preparation and imaging

Six milliliters of blood were collected via venipuncture from five healthy donors and equally divided between two 3 mL tubes of ethylenediaminetetraacetic acid (EDTA) (BD Medical 367856 Vacutainer®, United States) at the KAIST Clinic Pappalardo Center. One 3 mL sample of blood was tested using a CBC analyzer (XT-2000i, Sysmex Co., Kobe, Japan), and the other was tested using label-free tomographic imaging. Blood samples were stored in a refrigerator at 4°C and imaged using a tomographic microscope within 30 h after collection.

One micrometer of whole blood was loaded into the inlet of the developed device for imaging and filled into the entire microfluidic channel through capillary action. The blood samples in the EDTA tubes were placed at room temperature for imaging in a laboratory orbital shaker (SHO-2D, Daihan Scientific Co., Korea) to avoid blood coagulation. All experiments were approved by the Internal Review Board at KAIST.

### Quantification of RBC overlaps

We defined a metric that indicates the number of RBC layers based on the ratio of the sum of individual cell areas to the union of the cell areas as follows, to quantify the number of RBC layers using our scheme: α = Σ_*i*_, *A*_*i*_, /*U*_*i*_, *A*_*i*_, where *i*, *A,* and *U* are the indices of a cell, its area, and the union symbol, respectively. We manually segmented an individual cell and evaluated the metric for the three groups of devices fabricated with different channel heights to compute the area of each RBC because it was challenging to extract a cell mask for the highly overlapped RBCs using an automated segmentation algorithm.

## Results and discussions

### Fluid dynamics of capillary microfluidic device

Actuator-free capillary flow driven-microfluidic device was fabricated by utilizing glass substrate and SU-8 photoresist. As shown in Fig. 1f, one device is composed of 4 independent channels and has the total thickness less than 500 μm for the compatibility of 3D QPI imaging. The channel thickness of the device is about 2-3 μm, which makes it possible to observe RBC as a quasi-monolayer given that the thickness of the RBC is about 2 μm (Fig. 1g). The water contact angles of SiO_2_ and SU-8 constituting the surface of the channel are 69.6±2.7° and 94.1±1.9°, respectively (Fig. 1h). Since most of the channels are surrounded by SiO_2_, there is a flow due to negative capillary pressure that allows filling of the microchannel once the liquid is dropped at the entrance of the device without any external pump or actuators. The independence of actuators could facilitate usage of the device as a POC platform.

Figure 3 presents the dynamics of fluid in the microfluidic device using capillary forces with different heights. As our device relies on the negative capillary pressure arising from the interaction between the whole blood and the SiO_2_ channel, unlike many existing microfluidic devices with a flowing-rate-controllable pump, it is crucial to design the microchannel accurately to control its filling rate, which directly affects the blood testing time. The time required for the sample loading process to fill the channel was measured manually. Figure 3a presents the average filling rates of the three devices (device #1: 0.93±0.09 mm/s, device #2: 1.49±0.2 mm/s, and device #3: 2.67 mm/s). As expected, the filling rate is proportional to the device’s height because the negative capillary pressure-driven acceleration is inversely proportional to the channel height. However, the hydraulic resistance-driven acceleration is inversely proportional to the square of the channel height. As the liquid flow in the microchannel can be characterized by the channel dimensions and properties of the liquid, such as density, dynamic viscosity, and surface energy, the flow front position can be theoretically analyzed and compared with the experimental results (Fig. 3b). The theoretical model of the micro-scale fluidic channel assumes two dominant body forces induced by negative capillary pressure and hydraulic resistance, except for the gravitational effect. The experimental fluid flow agrees with the theoretical model, indicating that our microchannel devices were fabricated as designed.

**Figure 3.**
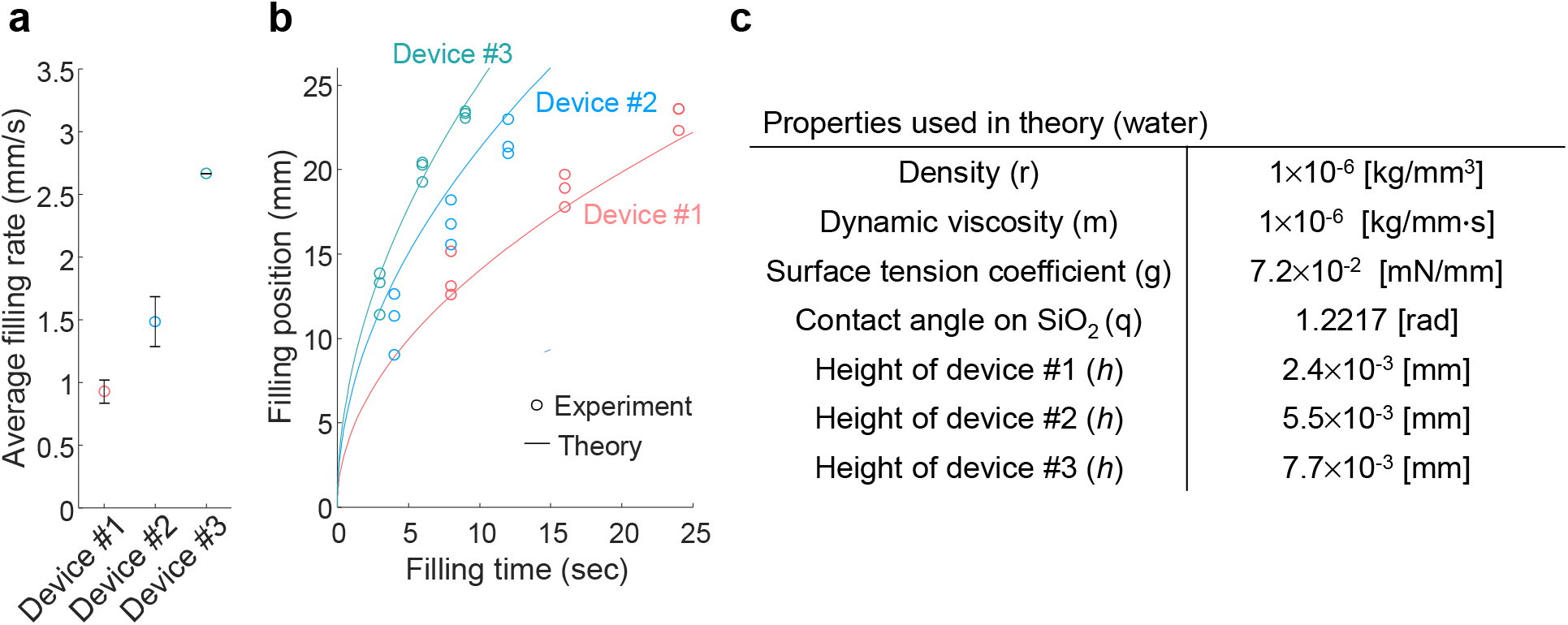
Rapid and actuator-free sample preparation using negative capillary pressure. (a) Average filling rate corresponding to the devices with different channel heights. Circles indicate average velocity, and error bars represent standard deviation. (b) Filling position-time plots for the three devices. Experimental results are compared to the theoretical model. The filling front position measurements, 8, 16, and 24 s, were measured for device #1; 4, 8, and 12 s were measured for device #2; 3, 6, and 9 s were measured for device #3. Circles and solid lines represent experimental values and theoretical curves, respectively.

When we apply the system to whole blood, as shear-thinning fluid, increased shear stress makes the viscosity of blood decrease, which contributes to the relieved hydraulic resistance. On the other hand, since blood contains RBCs, WBCs, platelets, plasma proteins, and etc, the sticky and viscous fluid would make the filling rate slower than water. In spite of these complex behaviors of blood, we speculate negligible difference on our platform because of their relatively short filling time and simple geometry.

### Optimal device dimension for monolayer RBC imaging

To identify the optimal dimension of our capillary microchannel device to enable rapid imaging of the monolayer of dilution-free RBCs, we fabricated devices with three different channel heights and quantified the number of RBC layers using a metric that calculates the ratio of the sum of individual cell areas to the union of the cell areas α (see Methods).

Figure 4a illustrates examples of RBC RI tomograms obtained using the three devices with the corresponding α values. Note that the tomograms were visualized using maximum intensity projection. While the RBCs spread out with analyzable density at the single-cell level as for device #1 (top row), the axially overlapped cells prepared in devices #2 and #3 (bottom row) obstructed the morphological characterization of individual RBCs, impeding biochemical and mechanical quantification. Additionally, such dense RBCs deteriorate the imaging performance of the diffraction tomography because the highly clustered cells may not just induce unexpected flows within the microchannel while being imaged but may also have large or abrupt phase delays that could degrade accurate tomographic reconstruction^44, 54^. Figure 4b shows the correlative scatter plot between α and the mean channel height of the three different devices. The mean and standard deviation of α for the three devices are 1.086 0.074, 1.327 0.066, and 1.728 ± 0.137, respectively, indicating that device #1 was used to characterize various properties of RBCs. We would also like to point out that it was challenging to fabricate SiO_2_ microchannels thinner than those used in device #1. Additionally, the flow of whole blood into such devices was also challenging as the top cover glasses occasionally warped or even collapsed due to low channel height.

**Figure 4.**
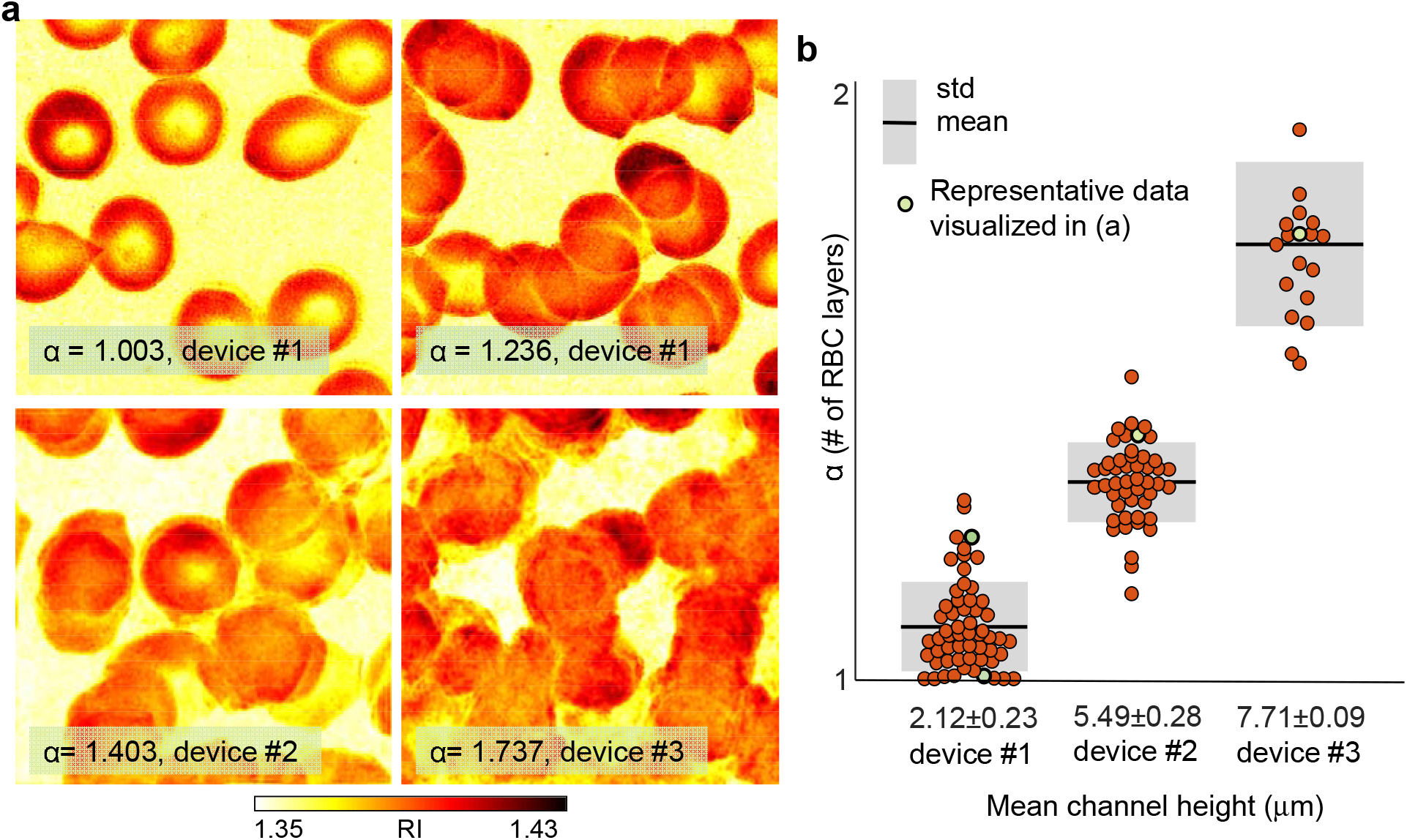
Quantification of RBC overlaps in different device dimensions. (a) Examples of RBC RI tomograms captured using the three devices. (b) Correlative scatter plot between the overlap metric, α, and device height. Green data points correspond to the RI tomograms of RBCs in (a).

### Morphological, biochemical, and mechanical quantification of RBCs

The morphological and biochemical properties of RBCs obtained from healthy donors were quantitatively investigated to validate the developed dilution- and label-free RBC testing framework. We reconstructed the 3D RI tomograms of RBCs in a wide field of view, using the proposed capillary microfluidic device that generates a monolayer of RBCs and obtained various properties from the RBC tomograms, including Hb concentration, Hb content, and cellular volume at the single-cell level (see also Methods). Figures 5a–c compare these results with those of the CBC test. The means and standard deviations of the Hb concentration, Hb content, and cellular volume were 34.55 ± 0.67 g/dL, 32.06 ± 5.90 pg, and 92.47 ± 17.20 fL, respectively. A total of 316 data points were collected. While our test results agree with the corresponding CBC results (orange dotted line), it is noteworthy that the Hb concentration results from our framework tended to be slightly lower than those from the CBC test. This is attributed to the underestimation of the RI values, which are linearly related to the Hb concentration, owing to the inaccessible information of the 3D transfer function^45^. We postulate that improved regularization algorithms that effectively compensate for the underestimated RI could yield a more accurate Hb concentration.

**Figure 5.**
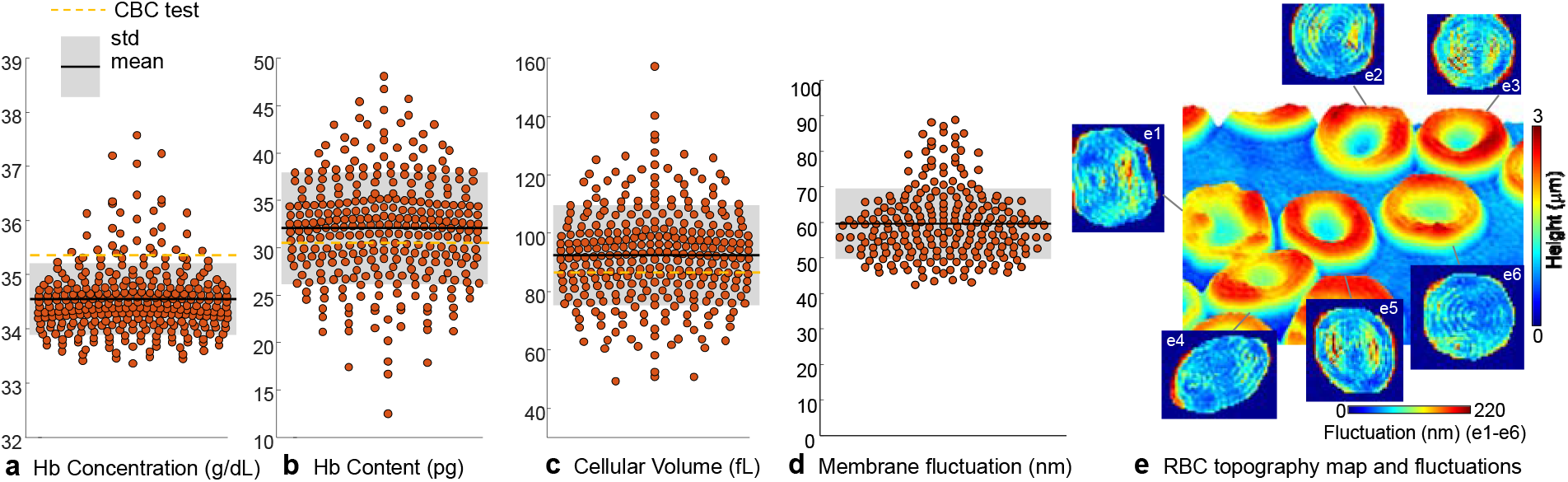
Quantitative parameters of RBCs. (a) Hb concentration. (b) Hb Content. (c) Cellular volume. (d-e) Membrane fluctuation and topography map. Yellow-dotted lines represent CBC results.

The time-series phase of RBCs was measured to study cell deformability, which is one of the important mechanical properties of RBCs, to validate the developed framework. As the topographical information of RBCs can be inverted directly from their phase map, our framework that retrieves the phase delay map of monolayered RBCs with normal illumination can be a powerful tool for probing the dynamic membrane fluctuation of RBCs. The analysis revealed that the dynamic membrane fluctuation of RBCs could be measured accurately using quantitative phase imaging techniques for diverse hematological applications^38, 55^. Figures 5(d-e) depict the membrane fluctuation of RBCs measured using our system with a representative topography map. The dynamic phase of RBCs obtained from one of the four healthy donors was measured, and the membrane fluctuation of each cell (mean ± std = 59.93 ± 9.91 nm, 232 RBCs) was quantified over a period of 2 s, which is in accordance with previous studies^56^. In this study, the membrane fluctuation is defined by spatially averaging the temporal standard deviation of each pixel’s height over an RBC area. A detailed calculation can be found in the sub-section titled “Quantification of RBC parameters.”

## Conclusions

This study reports a simple yet functional method for a completely reagent-free RBC analysis by exploiting capillary microfluidics and optical diffraction tomography. The 3D RI tomograms of RBCs are effectively obtained after the whole blood, without any treatment (e.g., dilutions, labeling) or additional microfluidic instruments (e.g., a pump, a mixer), is directly loaded into the microchannel, which generates a quasi-monolayer of RBCs. This enables the subsequent acquisition of their morphological (cell volume, 3D imaging) and biochemical parameters (Hb concentration and content) at the single-cell level. Our scheme also enables an RBC deformability study, which requires temporal stability inside the microchannel for time-series phase measurements. As the current study confirms that dilution-free RBC samples can be prepared within tens of seconds inside an ~2 μm-thick microfluidic device for optimal 3D imaging, it would be possible to further reduce the preparation time by fabricating a shorter channel. We believe that the developed approach can save significant time, costs, and labor involved in clinical settings, enabling point-of-care prognosis and diagnosis of various diseases. Several directions for future work remain to improve the developed approach. Based on the learnings from the microchannel developed for preparing RBC monolayers, we are attempting to fabricate capillary microchannel devices compatible with different blood cells, including white blood cells and platelets. Recently structured microchannels that exploit deterministic hydrodynamics for WBC separation could be excellent references for expanding the capabilities of our capillary-action-based device^57–59^. Additionally, advanced imaging processing approaches can also be employed^60^ or advanced cell segmentation algorithms for 3D imaging^61^ to achieve fully automated, high-throughput blood cell characterization. In this study, as partially overlapped RBCs inside the channel were captured in several FOVs, we manually segmented cell masks based on MIP images to quantify the RBC overlaps and extract their quantitative parameters. Improved algorithms that benefit from the full 3D information of RI tomograms may resolve the labor-intensive segmentation, augmenting the throughput of our approach. Furthermore, machine learning approaches can be readily combined with the present platform for rapid classifications of blood cell types and diagnosis of hematologic disorders^62, 63^. Finally, other imaging modalities could benefit from our capillary microfluidic device for low-cost, rapid characterization of biological samples^64, 65^.

## ASSOCIATED CONTENT

### Supporting Information

Theoretical model of capillary-driven sample loading. Properties used in the theoretical model.

## AUTHOR INFORMATION

### Author Contributions

After the project was initiated by Y.P., D.R and H.N conducted experiments and analyzed data under the supervision of J.S.J and Y.P. D.R prepared the draft manuscript and all authors revised it. ‡These authors contributed equally.

## ACKNOWLEDGMENTS

We thank Mahn Jae Lee and Young Seo Kim for assisting in sample preparation. We also thank Dr. Seungwoo Shin for his constructive comment on the quantitative RBC analysis.

## REFERENCES

1. Piagnerelli, M.; Boudjeltia, K. Z.; Vanhaeverbeek, M.; Vincent, J.-L., Red blood cell rheology in sepsis. Applied Physiology in Intensive Care Medicine 2009, 273–282.

2. Baskurt, O. K.; Gelmont, D.; Meiselman, H. J., Red blood cell deformability in sepsis. American journal of respiratory and critical care medicine 1998, 157 (2), 421–427.

3. Gulati, G. L.; Hyun, B. H., The automated CBC: a current perspective. Hematology/oncology clinics of North America 1994, 8 (4), 593–603.

4. Weatherby, D.; Ferguson, S., Blood chemistry and CBC analysis. Weatherby & Associates, LLC: 2002; Vol. 4.

5. DeNicola, D. B., Advances in hematology analyzers. Topics in companion animal medicine 2011, 26 (2), 52–61.

6. Winkelman, J. W.; Tanasijevic, M. J.; Zahniser, D. J., A novel automated slide-based technology for visualization, counting, and characterization of the formed elements of blood: A proof of concept study. Archives of pathology & laboratory medicine 2017, 141 (8), 1107–1112.

7. Atkins, C. G.; Buckley, K.; Blades, M. W.; Turner, R. F., Raman spectroscopy of blood and blood components. Applied spectroscopy 2017, 71 (5), 767–793.

8. Kim, D.; Oh, N.; Kim, K.; Lee, S.; Pack, C.-G.; Park, J.-H.; Park, Y., Label-free high-resolution 3-D imaging of gold nanoparticles inside live cells using optical diffraction tomography. Methods 2018, 136, 160–167.

9. Popescu, G.; Park, Y.; Choi, W.; Dasari, R. R.; Feld, M. S.; Badizadegan, K., Imaging red blood cell dynamics by quantitative phase microscopy. Blood Cells, Molecules, and Diseases 2008, 41 (1), 10–16.

10. Merola, F.; Memmolo, P.; Miccio, L.; Savoia, R.; Mugnano, M.; Fontana, A.; D’ippolito, G.; Sardo, A.; Iolascon, A.; Gambale, A., Tomographic flow cytometry by digital holography. Light: Science Applications 2017, 6 (4), e16241–e16241.

11. Memmolo, P.; Miccio, L.; Merola, F.; Gennari, O.; Netti, P. A.; Ferraro, P., 3D morphometry of red blood cells by digital holography. Cytometry part A 2014, 85 (12), 1030–1036.

12. Sinha, A.; Chu, T. T.; Dao, M.; Chandramohanadas, R., Single-cell evaluation of red blood cell bio-mechanical and nano-structural alterations upon chemically induced oxidative stress. Scientific reports 2015, 5 (1), 1–8.

13. Rinehart, M. T.; Park, H. S.; Walzer, K. A.; Chi, J.-T. A.; Wax, A., Hemoglobin consumption by P. falciparum in individual erythrocytes imaged via quantitative phase spectroscopy. Scientific reports 2016, 6 (1), 1–9.

14. Park, J.; Han, D. H.; Park, J.-K., Towards practical sample preparation in point-of-care testing: user-friendly microfluidic devices. Lab on a Chip 2020, 20 (7), 1191–1203.

15. Laxmi, V.; Tripathi, S.; Agrawal, A., Current Status of the Development of Blood-Based Point-of-Care Microdevices. Mechanical Sciences 2021, 169–196.

16. Park, H. S.; Eldridge, W. J.; Yang, W.-H.; Crose, M.; Ceballos, S.; Roback, J. D.; Chi, J.-T. A.; Wax, A., Quantitative phase imaging of erythrocytes under microfluidic constriction in a high refractive index medium reveals water content changes. Microsystems & nanoengineering 2019, 5 (1), 1–9.

17. Ramaswamy, B.; Yeh, Y.-T. T.; Zheng, S.-Y., Microfluidic device and system for point-of-care blood coagulation measurement based on electrical impedance sensing. Sensors and Actuators B: Chemical 2013, 180, 21–27.

18. Alazzam, A.; Stiharu, I.; Bhat, R.; Meguerditchian, A. N., Interdigitated comb-like electrodes for continuous separation of malignant cells from blood using dielectrophoresis. Electrophoresis 2011, 32 (11), 1327–1336.

19. Cooper, R. M.; Leslie, D. C.; Domansky, K.; Jain, A.; Yung, C.; Cho, M.; Workman, S.; Super, M.; Ingber, D. E., A microdevice for rapid optical detection of magnetically captured rare blood pathogens. Lab on a Chip 2014, 14 (1), 182–188.

20. Ruiz-Vega, G.; Arias-Alpízar, K.; de la Serna, E.; Borgheti-Cardoso, L. N.; Sulleiro, E.; Molina, I.; Fernàndez-Busquets, X.; Sánchez-Montalvá, A.; Del Campo, F. J.; Baldrich, E., Electrochemical POC device for fast malaria quantitative diagnosis in whole blood by using magnetic beads, Poly-HRP and microfluidic paper electrodes. Biosensors and Bioelectronics 2020, 150, 111925.

21. Reboud, J.; Bourquin, Y.; Wilson, R.; Pall, G. S.; Jiwaji, M.; Pitt, A. R.; Graham, A.; Waters, A. P.; Cooper, J. M., Shaping acoustic fields as a toolset for microfluidic manipulations in diagnostic technologies. Proceedings of the National Academy of Sciences 2012, 109 (38), 15162–15167.

22. Ohlsson, P.; Petersson, K.; Augustsson, P.; Laurell, T., Acoustic impedance matched buffers enable separation of bacteria from blood cells at high cell concentrations. Scientific reports 2018, 8 (1), 1–11.

23. Park, J.; Park, J.-K., Finger-actuated microfluidic device for the blood cross-matching test. Lab on a Chip 2018, 18 (8), 1215–1222.

24. Songjaroen, T.; Dungchai, W.; Chailapakul, O.; Henry, C. S.; Laiwattanapaisal, W., Blood separation on microfluidic paper-based analytical devices. Lab on a Chip 2012, 12 (18), 3392–3398.

25. Maria, M. S.; Rakesh, P.; Chandra, T.; Sen, A., Capillary flow-driven microfluidic device with wettability gradient and sedimentation effects for blood plasma separation. Scientific reports 2017, 7 (1), 1–12.

26. Zheng, Y.; Li, Q.; Hu, W.; Liao, J.; Zheng, G.; Su, M., Whole slide imaging of circulating tumor cells captured on a capillary microchannel device. Lab on a Chip 2019, 19 (22), 3796–3803.

27. Choi, Y.-S.; Im, M. K.; Lee, M. R.; Kim, C. S.; Lee, K.-H., Highly sensitive enclosed multilayer paper-based microfluidic sensor for quantifying proline in plants. Analytica chimica acta 2020, 1105, 169–177.

28. Yeh, E.-C.; Fu, C.-C.; Hu, L.; Thakur, R.; Feng, J.; Lee, L. P., Self-powered integrated microfluidic point-of-care low-cost enabling (SIMPLE) chip. Science advances 2017, 3 (3), e1501645.

29. Dimov, I. K.; Basabe-Desmonts, L.; Garcia-Cordero, J. L.; Ross, B. M.; Ricco, A. J.; Lee, L. P., Stand-alone self-powered integrated microfluidic blood analysis system (SIMBAS). Lab on a Chip 2011, 11 (5), 845–850.

30. Zhu, Z.; Guan, Z.; Jia, S.; Lei, Z.; Lin, S.; Zhang, H.; Ma, Y.; Tian, Z. Q.; Yang, C. J., Au@ Pt nanoparticle encapsulated target-responsive hydrogel with volumetric bar-chart chip readout for quantitative point-of-care testing. Angewandte Chemie International Edition 2014, 53 (46), 12503–12507.

31. Song, Y.; Zhang, Y.; Bernard, P. E.; Reuben, J. M.; Ueno, N. T.; Arlinghaus, R. B.; Zu, Y.; Qin, L., Multiplexed volumetric bar-chart chip for point-of-care diagnostics. Nature communications 2012, 3 (1), 1–9.

32. Jong, W.; Kuo, T.; Ho, S.; Chiu, H.; Peng, S., Flows in rectangular microchannels driven by capillary force and gravity. International communications in heat and mass transfer 2007, 34 (2), 186–196.

33. Chakraborty, S., Dynamics of capillary flow of blood into a microfluidic channel. Lab on a Chip 2005, 5 (4), 421–430.

34. Kim, Y.; Shim, H.; Kim, K.; Park, H.; Jang, S.; Park, Y., Profiling individual human red blood cells using common-path diffraction optical tomography. Scientific reports 2014, 4 (1), 1–7.

35. Park, H.; Lee, S.; Ji, M.; Kim, K.; Son, Y.; Jang, S.; Park, Y., Measuring cell surface area and deformability of individual human red blood cells over blood storage using quantitative phase imaging. Scientific reports 2016, 6 (1), 1–10.

36. Brochard, F.; Lennon, J., Frequency spectrum of the flicker phenomenon in erythrocytes. Journal de Physique 1975, 36 (11), 1035–1047.

37. Bao, G.; Suresh, S., Cell and molecular mechanics of biological materials. Nature materials 2003, 2 (11), 715–725.

38. Park, Y.; Diez-Silva, M.; Popescu, G.; Lykotrafitis, G.; Choi, W.; Feld, M. S.; Suresh, S., Refractive index maps and membrane dynamics of human red blood cells parasitized by Plasmodium falciparum. Proceedings of the National Academy of Sciences 2008, 105 (37), 13730–13735.

39. Chandramohanadas, R.; Park, Y.; Lui, L.; Li, A.; Quinn, D.; Liew, K.; Diez-Silva, M.; Sung, Y.; Dao, M.; Lim, C. T., Biophysics of malarial parasite exit from infected erythrocytes. PloS one 2011, 6 (6), e20869.

40. Shin, S.; Kim, K.; Kim, T.; Yoon, J.; Hong, K.; Park, J.; Park, Y. In Optical diffraction tomography using a digital micromirror device for stable measurements of 4D refractive index tomography of cells, Quantitative Phase Imaging II, International Society for Optics and Photonics: 2016; p 971814.

41. Goldstein, R. M.; Zebker, H. A.; Werner, C. L., Satellite Radar Interferometry - Two-Dimensional Phase Unwrapping. Radio Sci 1988, 23 (4), 713–720.

42. Wolf, E., Three-dimensional structure determination of semi-transparent objects from holographic data. Optics communications 1969, 1 (4), 153–156.

43. Chen, B.; Stamnes, J. J., Validity of diffraction tomography based on the first Born and the first Rytov approximations. Applied optics 1998, 37 (14), 2996–3006.

44. Habashy, T. M.; Groom, R. W.; Spies, B. R., Beyond the Born and Rytov approximations: A nonlinear approach to electromagnetic scattering. Journal of Geophysical Research: Solid Earth 1993, 98 (B2), 1759–1775.

45. Lim, J.; Lee, K.; Jin, K. H.; Shin, S.; Lee, S.; Park, Y.; Ye, J. C., Comparative study of iterative reconstruction algorithms for missing cone problems in optical diffraction tomography. Optics express 2015, 23 (13), 16933–16948.

46. Lauer, V., New approach to optical diffraction tomography yielding a vector equation of diffraction tomography and a novel tomographic microscope. Journal of Microscopy 2002, 205 (2), 165–176.

47. Park, C.; Shin, S.; Park, Y., Generalized quantification of three-dimensional resolution in optical diffraction tomography using the projection of maximal spatial bandwidths. JOSA A 2018, 35 (11), 1891–1898.

48. Kim, K.; Yoon, J.; Shin, S.; Lee, S.; Yang, S.-A.; Park, Y., Optical diffraction tomography techniques for the study of cell pathophysiology. Journal of Biomedical Photonics & Engineering 2016, 2 (2).

49. Kim, K.; Yoon, H.; Diez-Silva, M.; Dao, M.; Dasari, R. R.; Park, Y., High-resolution three-dimensional imaging of red blood cells parasitized by Plasmodium falciparum and in situ hemozoin crystals using optical diffraction tomography. Journal of biomedical optics 2013, 19 (1), 011005.

50. Barer, R., Refractometry and interferometry of living cells. JOSA 1957, 47 (6), 545–556.

51. Popescu, G.; Park, Y.; Lue, N.; Best-Popescu, C.; Deflores, L.; Dasari, R. R.; Feld, M. S.; Badizadegan, K., Optical imaging of cell mass and growth dynamics. American Journal of Physiology-Cell Physiology 2008, 295 (2), C538–C544.

52. Park, Y.; Best, C. A.; Auth, T.; Gov, N. S.; Safran, S. A.; Popescu, G.; Suresh, S.; Feld, M. S., Metabolic remodeling of the human red blood cell membrane. Proceedings of the National Academy of Sciences 2010, 107 (4), 1289–1294.

53. Shaked, N. T.; Satterwhite, L. L.; Truskey, G. A.; Wax, A. P.; Telen, M. J., Quantitative microscopy and nanoscopy of sickle red blood cells performed by wide field digital interferometry. Journal of biomedical optics 2011, 16 (3), 030506.

54. Kak, A. C.; Slaney, M., Principles of computerized tomographic imaging. Society for Industrial and Applied Mathematics: 2001.

55. Park, Y.; Best, C. A.; Badizadegan, K.; Dasari, R. R.; Feld, M. S.; Kuriabova, T.; Henle, M. L.; Levine, A. J.; Popescu, G., Measurement of red blood cell mechanics during morphological changes. Proceedings of the National Academy of Sciences 2010, 107 (15), 6731–6736.

56. Lee, S.; Park, H.; Kim, K.; Sohn, Y.; Jang, S.; Park, Y., Refractive index tomograms and dynamic membrane fluctuations of red blood cells from patients with diabetes mellitus. Sci Rep 2017, 7 (1), 1039.

57. Davis, J. A.; Inglis, D. W.; Morton, K. J.; Lawrence, D. A.; Huang, L. R.; Chou, S. Y.; Sturm, J. C.; Austin, R. H., Deterministic hydrodynamics: Taking blood apart. Proceedings of the National Academy of Sciences 2006, 103 (40), 14779–14784.

58. Choi, S.; Song, S.; Choi, C.; Park, J.-K., Continuous blood cell separation by hydrophoretic filtration. Lab on a Chip 2007, 7 (11), 1532–1538.

59. Kim, B.; Kim, K. H.; Chang, Y.; Shin, S.; Shin, E.-C.; Choi, S., One-Step Microfluidic Purification of White Blood Cells from Whole Blood for Immunophenotyping. Analytical Chemistry 2019, 91 (20), 13230–13236.

60. Lee, J.; Kim, H.; Cho, H.; Jo, Y.; Song, Y.; Ahn, D.; Lee, K.; Park, Y.; Ye, S., Deep-Learning-Based Label-Free Segmentation of Cell Nuclei in Time-Lapse Refractive Index Tomograms. IEEE Access 2019, 7, 83449–83460.

61. Carvalho, L. E.; Sobieranski, A. C.; von Wangenheim, A., 3D Segmentation Algorithms for Computerized Tomographic Imaging: a Systematic Literature Review. Journal of Digital Imaging 2018, 31 (6), 799–850.

62. Kim, G.; Jo, Y.; Cho, H.; Min, H.-s.; Park, Y., Learning-based screening of hematologic disorders using quantitative phase imaging of individual red blood cells. Biosensors and Bioelectronics 2019, 123, 69–76.

63. Ryu, D.; Kim, J.; Lim, D.; Min, H.-S.; You, I.; Cho, D.; Park, Y., Label-free bone marrow white blood cell classification using refractive index tomograms and deep learning. bioRxiv 2020.

64. Myers, F. B.; Lee, L. P., Innovations in optical microfluidic technologies for point-of-care diagnostics. Lab on a Chip 2008, 8 (12), 2015–2031.

65. Breslauer, D. N.; Lee, P. J.; Lee, L. P., Microfluidics-based systems biology. Molecular Biosystems 2006, 2 (2), 97–112.

